# Control of β-glucan exposure by the endo-1,3-glucanase Eng1 in *Candida albicans* modulates virulence

**DOI:** 10.1101/2020.09.07.285791

**Authors:** Mengli Yang, Norma V. Solis, Michaela Marshall, Rachel Garleb, Tingting Zhou, Daidong Wang, Marc Swidergall, Eric Pearlman, Scott G. Filler, Haoping Liu

## Abstract

*Candida albicans* is a major cause of invasive candidiasis, which has a high mortality rate. The hyphal form of *C. albicans* is virulent and activates the host innate immune response, while the yeast form is hypovirulent and less immunogenic. The innate immune response is critical for host defense, but overactivation can cause tissue damage and sepsis. The innate immune response can be triggered when the C-type lectin receptor Dectin-1 recognizes β-glucans, which is protected by the outer mannan layer of the cell wall on *C. albicans*. Here, we demonstrate that there is low level of Dectin-1 binding at the septum of yeast cells, but high level of Dectin-1 binding over the entire surface of hyphae. We find that β-glucan masking in yeast is controlled by two highly expressed yeast proteins, the endo-1,3-β-glucanase Eng1 and the Yeast Wall Protein Ywp1. An *eng1* deletion mutant shows enhanced Dectin-1 binding at the septa, while an *eng1 ywp1* double mutant, but not an *ywp1* single mutant, shows strong overall Dectin-1 binding. Thus, Eng1-mediated β-glucan trimming and Ywp1-mediated β-glucan masking are two parallel mechanisms utilized by *C. albicans* yeast to minimize recognition by Dectin-1. In the model of disseminated candidiasis, mice infected with the *eng1* deletion mutant showed delayed mortality with an increased renal immune response in males compared to mice infected with the wild-type strain, but earlier mortality with a higher renal immune response in females. Using the *eng1* mutant that is specifically defective in β-glucan masking in yeast, this study demonstrates that the level of β-glucan exposure is important for modulating the balance between immune protection and immunopathogenesis.

**Abstract Importance:** *Candida albicans* is a major opportunistic fungal pathogen of humans. Systemic Candidiasis has high mortality rates. *C. albicans* is also a constituent of the human microbiome and found in gastrointestinal and genitourinary tracts of most healthy individuals. *C. albicans* is able to switch reversibly between yeast and hyphae in response to environmental cues. The hyphal form is virulent, while the yeast form is hypovirulent and less immunogenic. This study demonstrates that β-glucan exposure in yeast is protected by two highly expressed yeast proteins, the endo-1,3-β-glucanase Eng1 and the Yeast Wall Protein Ywp1. Eng1-mediated β-glucan trimming and Ywp1-mediated β-glucan masking are two parallel mechanisms utilized by *C. albicans* yeast to minimize recognition by the host C-type lectin receptor Dectin-1. The *eng1* mutant triggers a higher immune response and leads to earlier mortality compared to the wild-type strain. Thus, β-glucan masking in yeast keeps yeast cells less immunogenic and hypovirulent.

## Introduction

Systemic candidiasis, mainly caused by the opportunistic yeast *Candida albicans*, is the fourth leading cause of nosocomial bloodstream infections. The mortality of patients with invasive candidiasis exceeds 40% even with antifungal treatment (1,2). The innate immune response is critical for the host defense against invasive candidiasis, but it can also be detrimental by causing tissue damage and sepsis (3–5). Thus, it is critically important to understand how *C. albicans* regulates its recognition by host immune cells to modulate the host immune response.

The host immune response to candidal infection is initiated when pattern-recognition receptors (PRR) on innate immune cells recognize *Candida* cell wall carbohydrates, which serve as pathogen-associated molecular patterns (PAMPs). A major PRR for fungi such as *C. albicans* is Dectin-1, a C-type lectin-receptor that can recognize β-1,3-glucan on the fungal cell wall and is required for the host defense against hematogenously disseminated candidiasis (6,7). The *C. albicans* cell wall consists of an outer layer of mannosylated proteins and an inner layer of mostly β-1,3-glucan and underlying chitin (8). Under normal conditions, β-1,3-glucan is masked from Dectin-1 detection by the outer mannan layer (8). β-glucan exposure is affected by many factors, including growth conditions (e.g. Lactate (9), hypoxia (10), iron depletion (11)), mutations in mannan synthesis pathways (12), antifungal treatment (13,14), and different *C. albicans* strain backgrounds (15), all in support of the current model that β-glucan is masked by the outer mannan layer (8). Different from the mannan masking model, exoglucanase Xog1 is recently shown to reduce β-glucan exposure in lactate (16). *Histoplasma capsulatum* secretes an endo-1,3-β-glucanase (Eng1) to trim down exposed β-glucan from the cell surface to prevent Dectin-1 recognition (17). Whether a similar mechanism is used by *Candida* species is not clear.

The ability to transition between yeast and hyphae is critical for the virulence of *C. albicans* (18). Hyphae are considered to be the virulent form of *C. albicans* that facilitates host invasion and triggers host immune response activation, while yeast are regarded as hypovirulent and less immunogenic (18,19). Several studies have indicated differential cytokine responses of innate immune cells to yeast and hyphae. Inflammasome activation is preferentially induced by *Candida* hyphae, and this reflects an important mechanism of discrimination between colonization and invasion (8). The increased inflammasome activation/IL-1β production in response to hyphae is partially mediated by Dectin-1, suggesting an increased recognition of β-glucans in hyphae, which is thought to be caused by the shorter and less abundant mannan fibrils on the surface of hyphae (20). The molecular mechanisms for the proposed regulation of mannan layer thickness or β-glucan exposure in yeast and hyphae are elusive. Whether β-glucan on *C. albicans* hyphae is more exposed and better recognized by Dectin-1 remains controversial (21–23). This is further complicated by the fact that most studies on β-glucan masking used inactivated yeast cells to study immune response, as *C. albicans* yeast cells develop into hyphae upon exposure to myeloid cells (macrophage and dendritic cells). Here, we show that hyphae are better recognized by Dectin-1 than yeast cells. Further, we provide molecular mechanism and regulators for yeast-specific β-glucan masking. The importance of β-glucan masking in yeast for *C. albicans* virulence and immunopathogenesis is examined in the model of hematogenously disseminated candidiasis.

## Results

### β-glucan is masked in yeast but not hyphae

Gantner *et al*. reported that β-glucan exposure was detectable only on yeast cells at the bud scar, but not on hyphae (21). In contrast, high β-glucan exposure was observed in hyphae, but not in yeast, when cells were treated with a sub-inhibitory concentration of caspofungin (14). Dectin-1 binding on hyphae was also observed during infection (14). Two recent investigations also observed Dectin-1 binding to hyphae, although no direct comparison were made between yeast and hyphae (22,23). Here, we directly compared Dectin-1 binding between yeast and hyphae, using Dectin-1-Fc and a fluorescein-conjugated secondary antibody. Yeast cells showed Dectin-1 binding only at the bud scar, as reported previously(21) (Fig 1a). In contrast, Dectin-1 binding was observed on the cell wall of germ tubes, but not on the basal yeast cells (Fig 1a). ImageJ analysis showed that hyphae had higher levels of Dectin-1-Fc binding than yeast (Fig 1a, right panel). To examine if the higher β-glucan exposure in hyphae was linked to higher immune activation, we infected bone marrow-derived macrophages (BMDM) with live yeast-locked *flo8* cells and live wild-type yeast cells, which rapidly developed into hyphae during infection. The wild-type hyphae induced significantly higher level of TNFα than the *flo8* yeast cells (Fig 1b). The cytokine response to yeast cells was largely Dectin-1 dependent, but only partially Dectin-1 dependent in response to hyphae, indicating that the BMDMs also responded to other PAMPs on hyphae. We further showed that induction of TNFα by hyphae was largely blocked by the SYK inhibitor R406, suggesting that immune activation by hyphae is mediated through C-type lectin receptors (Fig 1c). The higher immune response triggered by hyphae could be due to larger surface area, and therefore, more PAMPs. How β-glucan masking is differentially regulated in yeast and hyphae is not clear, but β-glucan is the major PAMP for immune activation in yeast.

**Figure 1.**
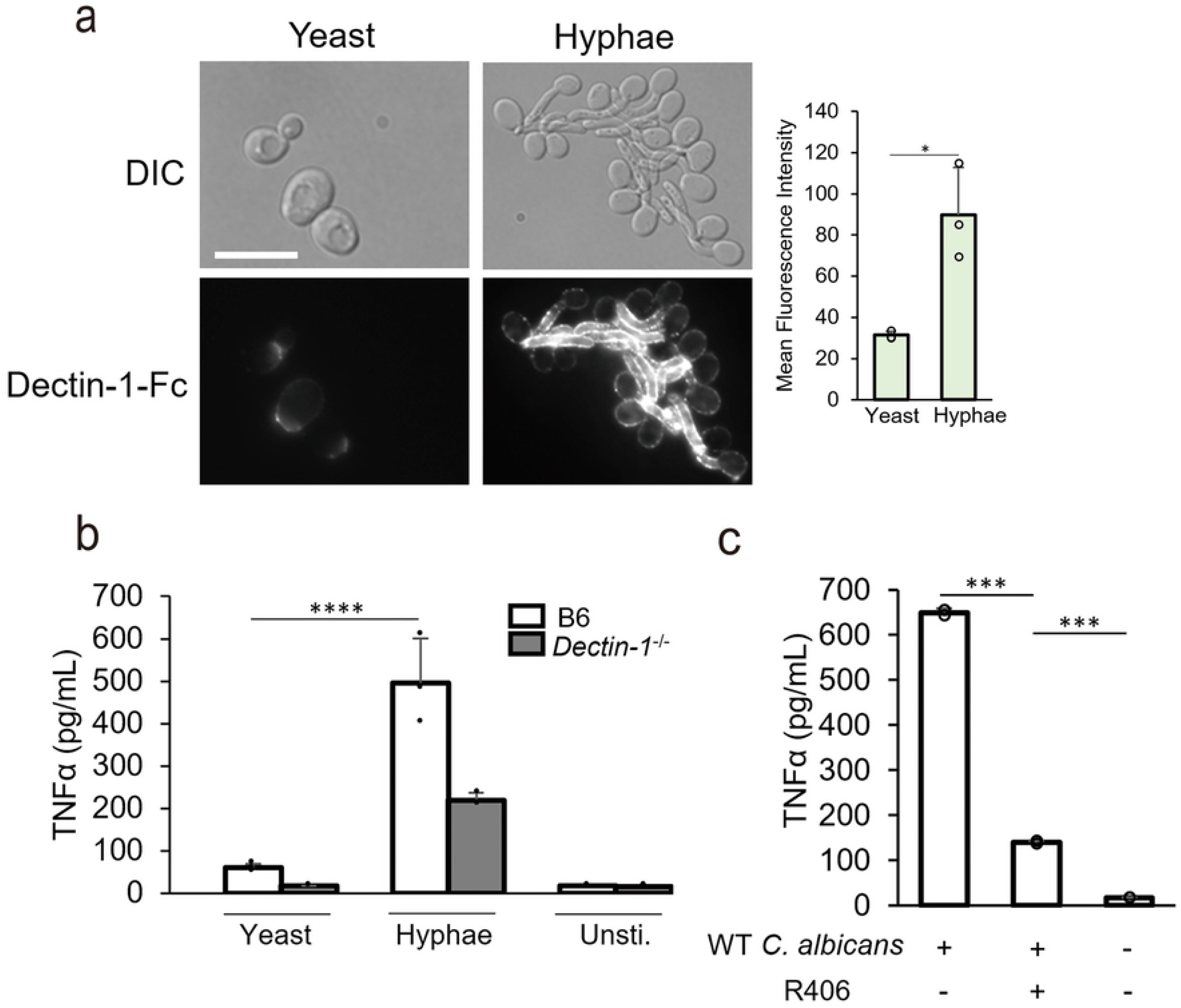
β-glucan is masked in yeast and exposed in hyphae. (a) Representative images of *C. albicans* cells stained with Dectin-1-Fc and secondary antibody conjugated to FITC. Yeast cells were cultured in YPD at 30 degrees for 6 hours. Hyphae were induced in YPD at 37 degrees for 1 hour. The scale bar represents 10μM. Mean fluorescence intensities were quantitated by ImageJ. (b) The levels of TNFα in the supernatant of BMDM stimulated with *flo8* and WT live yeast by a MOI of 1:1. (c) The levels of TNFα in the supernatant of BMDM stimulated with live hyphae with or without treatment of Syk inhibitor R406. *p* Values were calculated using one-way ANOVA with Tukey post hoc analysis (\*\*\*\**p*<0.0001; *** *p*<0.001)

### Control of β-glucan exposure by the endo-1,3-glucanase Eng1 in yeast

The higher β-glucan exposure in hyphae relative to yeast could be attributed to faster growth of hyphal cell wall, or a thinner outer layer of mannan (20). It is also possible that there are yeast-specific mechanisms of β-glucan protection. Instead of masking of β-glucan as occurs in *C. albicans, Histoplasma capsulatum* produces an endo-1,3-β-glucanase (Eng1) to digest exposed β-glucan on the cell wall (17). *C. albicans* has an Eng1 ortholog (24) that is regulated by Ace2 (25), a transcription factor required for septum destruction after cytokinesis as in *Saccharomyces cerevisiae* (26). At early G1 phase in yeast, *ENG1* is expressed and secreted from the daughter cell to degrade β-glucan in the septum, leaving a bud scar on the mother side (Fig 2a) (26). *ENG1* expression is downregulated in hyphae (Fig 2b), as Efg1 phosphorylation by hypha-specific Cdk1-Hgc1 represses Ace2-regulated genes (Fig 2a) (27), leading to the formation of chains of cells with increased β-glucan exposure.

**Figure 2.**
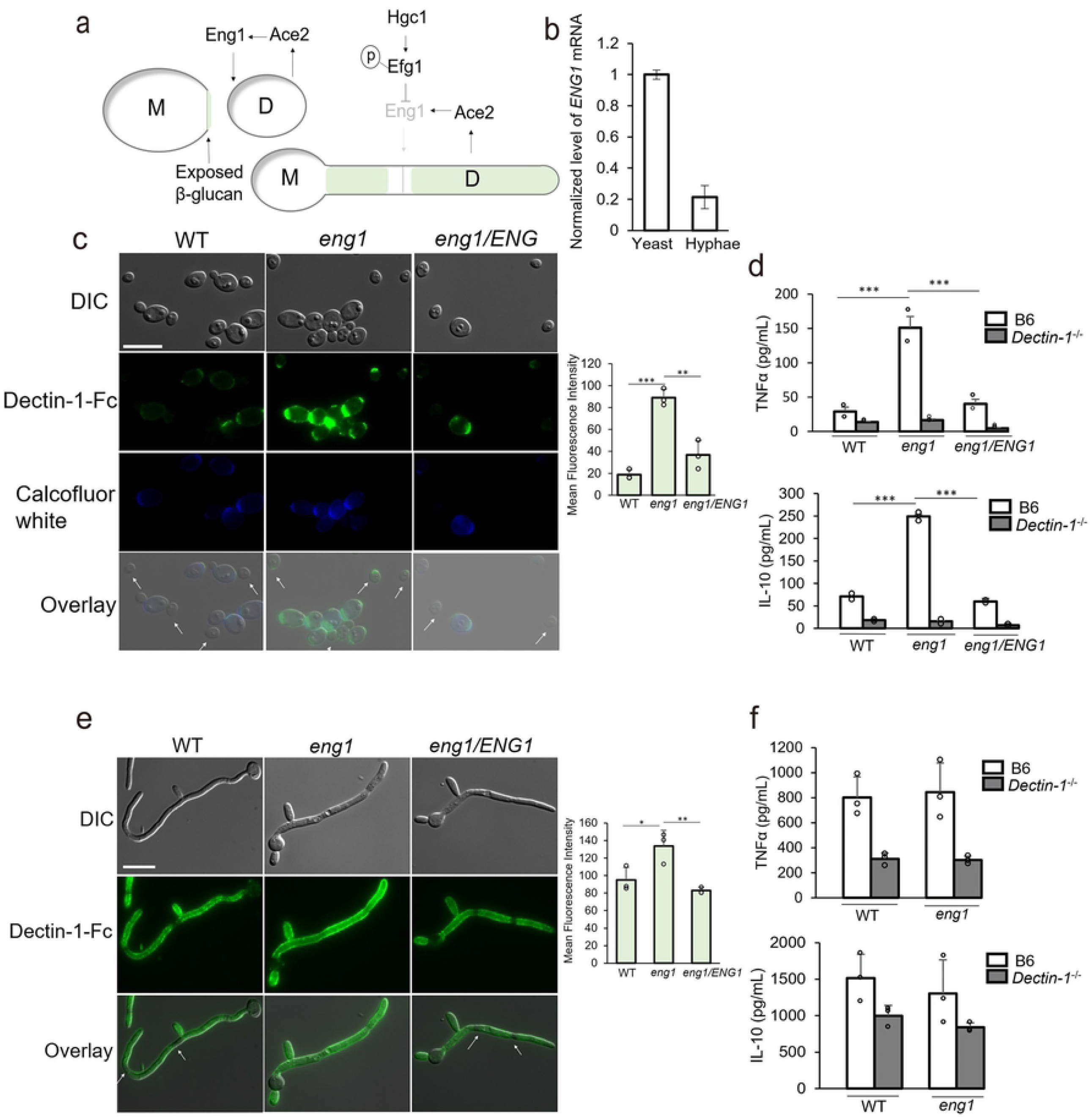
Eng1 reduces β-glucan exposure in yeast. (a) Regulation of Eng1 expression in yeast and hyphae. (b) Expression levels of *ENG1* mRNA in yeast or hyphae quantitated by qPCR. Yeast cells were cultured in YPD at 30 degrees for 6 hours. Hyphae were cultured in YPD at 37 degrees for 1 hour. (c) Representative images of *C. albicans* cells stained with Dectin-1-Fc and secondary antibody conjugated to FITC. Yeast cells were cultured in YPD for 6 hours. The scale bar represents 10μM. Mean fluorescence intensities were quantitated by ImageJ. (d) The levels of TNFα and IL-10 in the supernatant of BMDM stimulated with fixed *C. albicans* yeast form by a MOI of 1:3. Asterisks (***) indicate *p*<0.001 based on one-way ANOVA with Tukey post hoc analysis. (e) Representative images of Hyphae stained with Dectin-1-Fc and secondary antibody conjugated to FITC. Hyphae were induced in SC with 2% N-acetylglucosamine for 5 hours. Mean fluorescence intensities were quantitated by ImageJ. The scale bar represents 10μM. (f) The levels of TNFα and IL-10 in the supernatant of BMDM stimulated with WT or eng1 live yeast by a MOI of 1:1. Asterisks (**) indicate *p*<0.01 based on one-way ANOVA with Tukey post hoc analysis.

To determine if Eng1 is involved in trimming exposed β-glucan, we constructed an *eng1* deletion by CRISPR/Cas9 (28). The *eng1* mutant showed an increase in cell chain formation and enhanced Dectin-1 binding at septa (Fig 2c). In WT and *ENG1* complemented *eng1* cells, Dectin-1 binding was only seen at the bud scar of mother cells, as indicated by chitin staining with calcofluor white, but not daughter cells (white arrow). In *eng1* cells, Dectin-1 binding was also detectable in daughter cells at regions where chitin was not exposed (white arrows). Dectin-1 binding of *eng1* yeast was 3-fold higher than WT yeast (Fig 2c). Consistent with their increased Dectin-1 recognition, fixed *eng1* yeast cells induced higher levels of TNFα and IL-10 in BMDM compared to the fixed yeast of the WT or *ENG1* complemented strain (Fig 2d). Cytokine production was greatly reduced in Dectin-1 deficient BMDM, indicating that the immune response to the *eng1* mutant in yeast cells is dependent on Dectin-1.

In hyphae, the *eng1* mutant showed uniform Dectin-1-Fc staining along the cell walls even at mother-daughter junction sites, which were masked in wild-type hyphae (Fig 2e, white arrows). The overall level of Dectin-1-Fc staining was slightly higher in *eng1* hyphae than in WT hyphae, but *eng1* hyphae induced WT levels of TNFα and IL-10 production by BMDMs (Fig 2f). Although Dectin-1^-/-^ BMDMs secreted less TNFα and IL-10 than WT cells, they still produced a significant amount of both cytokines. Therefore, Eng1 plays a limited role in immune activation by hyphae and the BMDM response to hyphae is only partially dependent on Dectin-1.

### Co-regulation of β-glucan masking in yeast by Eng1 and the Yeast Wall Protein Ywp1

While Eng1 contributes to the trimming of β-glucan mainly on septa, the overall level of Dectin-1 binding to *eng1* yeast cells remained low (Fig 2c), indicating the existence of an additional yeast-specific mechanism for β-glucan masking. The Yeast Wall Protein (Ywp1), an anti-adhesin that is expressed highly in yeast cells, is implicated in blocking the accessibility to both anti-Ywp1 and anti-β-glucan antibodies (22). We constructed a *ywp1* single deletion mutant and an *eng1 ywp1* double deletion mutant using CRISPR/Cas9. The *ywp1* mutant showed slightly increased overall Dectin-1 binding and tended to form aggregates (Fig 3a). In comparison, the *eng1 ywp1* double mutant showed strong Dectin-1 binding at the septa and over the entire surface of the yeast cells (Fig 3a). The *ywp1 eng1* double mutant also induced much higher levels of TNFα and IL-10 secretion by BMDMs than the *ywp1* or *eng1* single deletion mutants (Fig 3b). Thus, deleting *ENG1* revealed the function of Ywp1 in β-glucan masking. Similarly, the β-glucanase activity of Eng1 on the entire cell wall was revealed in the absence of Ywp1 masking. Although cytokine production by BMDM in response to the single mutants was significantly lower in Dectin-1 deficient macrophages, there was only partial reduction in Dectin-1^-/-^ cells stimulated with the *ywp1 eng1* double mutant (Fig 3b). This could be due to frustrated phagocytosis of macrophages induced by large cell aggregates (29), which amplified the signals mediated through other C-type lectin receptors. Alternatively, the absence of both Ywp1 and Eng1 could result in the exposure of additional PAMPs other than β-glucan. Our finding of synergistic regulation of β-glucan masking by two highly expressed yeast proteins directly links the dimorphic regulation of β-glucan exposure to the hyphal transcriptional program (30).

**Figure 3.**
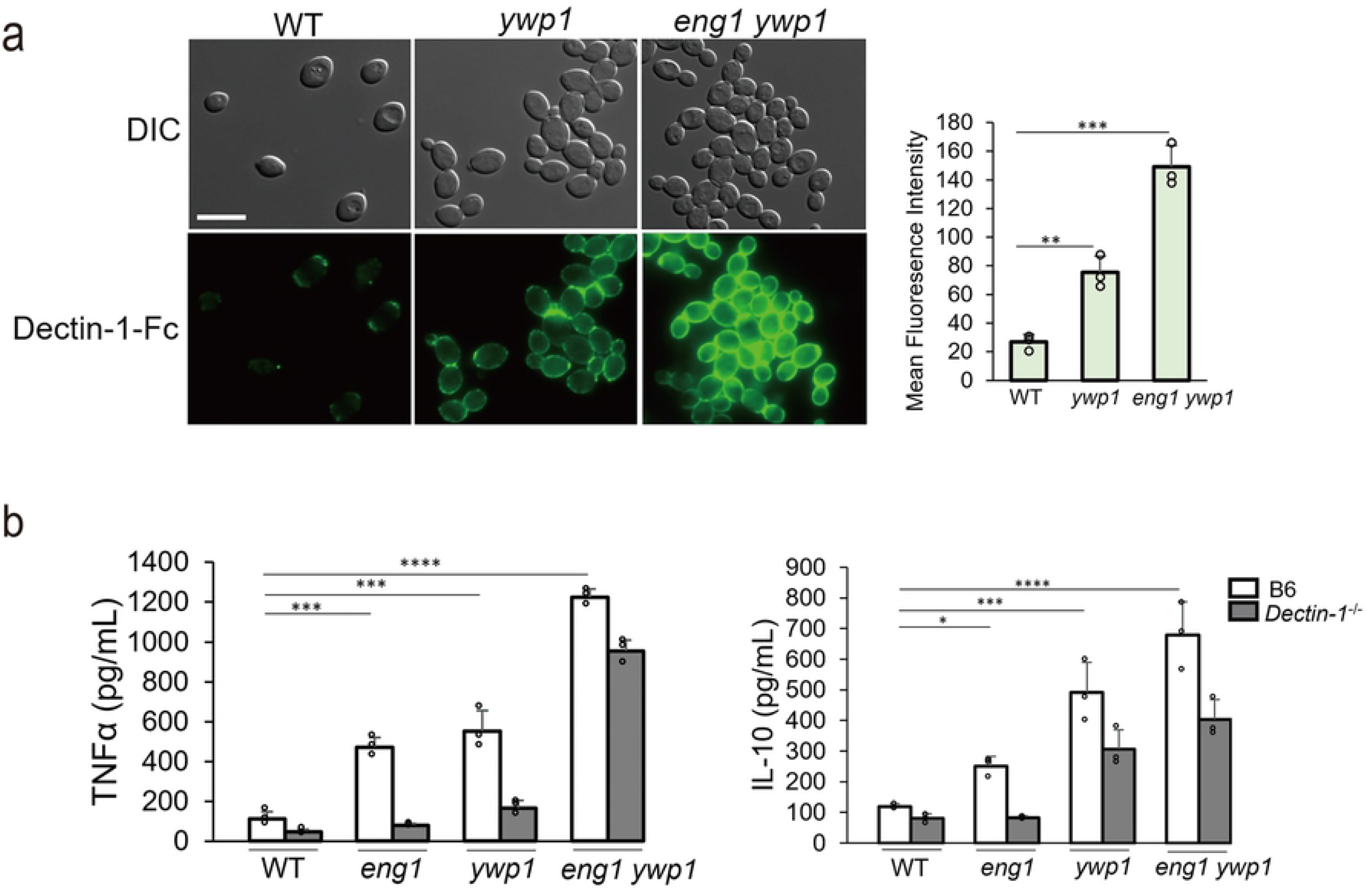
Regulation of β-glucan exposure in yeast by Eng1 and Ywp1. (a) Representative images of *C. albicans* yeast cells stained with Dectin-1-Fc and secondary antibody conjugated to FITC. Mean fluorescence intensities were quantitated by ImageJ. The scale bar represents 10μM. (b) The levels of TNFα or IL-10 in the supernatant of BMDM stimulated with fixed yeast cells by a MOI of 1:3. *p* Values were calculated using ANOVA with Tukey post hoc analysis (\*\*\*\**p*<0.0001; \*\*\**p*<0.001, \*\**p*<0.01, \**p*<0.05).

### *ENG1* expression in yeast cells is highly regulated

We also investigated whether *ENG1* expression is altered in response to treatments and conditions known to affect cell separation. Anti-fungal drugs fluconazole and caspofungin treated cells have been shown to increase cell chain formation, indicating a repression of Ace2-target genes (13,31). Indeed, *ENG1* transcript levels were lower in fluconazole or caspofungin treated cells (Fig 4a). The drug-treated cells induced higher levels of TNFα in BMDMs in a Dectin-1 dependent manner (Fig 4b), indicating an increase in Dectin-1 recognition. To determine if the higher Dectin-1 recognition is due to the down-regulation of Eng1, fixed WT, *eng1* and drug treated WT yeast cells were incubated overnight with the supernatant of WT culture, which should contain the secreted Eng1 enzyme. The supernatant from *eng1* cells was used as a no-Eng1 control. The *eng1* cells treated with WT supernatant had reduced Dectin-1-Fc binding as compared to the cells treated with *eng1* supernatant (Fig 4c). By contrast, the WT supernatant had no effect on WT cells. These results confirmed the β-glucan trimming ability of the Eng1 enzyme in the WT supernatant, even though it was not able to completely remove β-glucan at septum. When the cells were treated with fluconazole or caspofungin and incubated with the *eng1* supernatant, they bound more Dectin-1-Fc than the WT cells (Fig 4c). Fluconazole treated yeast cells had enhanced Dectin-1 binding at the septa between mother and daughter cells, similar to the *eng1* mutant. Caspofungin-treated yeast cells had an increase in overall Dectin-1-Fc binding. When the cells were treated with either antifungal agent, incubation with WT supernatant largely reduced the Dectin-1 binding, indicating that supplemental Eng1 enzyme was able to remove exposed β-glucan on the cell wall (Fig 4c). This suggested that reduced Eng1 expression contributes to the increased β-glucan exposure when *C. albicans* cells are exposed to fluconazole or caspofungin.

**Figure 4.**
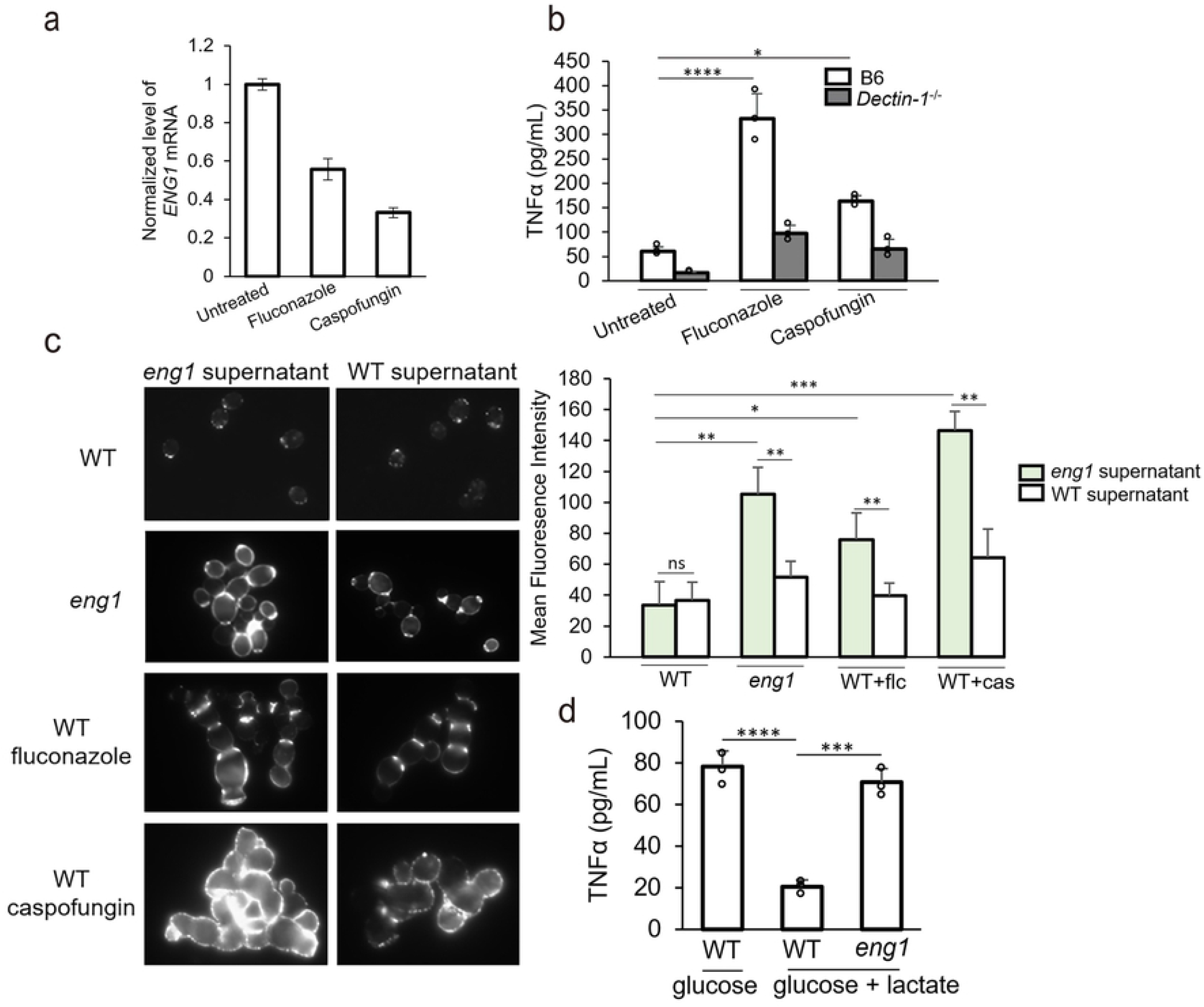
Down-regulation of *ENG1* during cell chain formation is associated with increased β-glucan exposure (a) Transcript levels of *ENG1* in fluconazole/caspofungin-treated or untreated cells. (b) The levels of TNFα in the supernatant of BMDM stimulated with fixed yeast cells by a MOI of 1:3. Cells were cultured in YPD with 10ug/mL fluconazole for overnight or 0.06ug/mL Caspofungin for 3 hours. (c) Representative images of yeast form cells stained with Dectin-1-Fc and secondary antibody conjugated to FITC. WT and *eng1* supernatant were collected and filtered from saturated overnight culture of WT or *eng1* yeast cells. Mean fluorescence intensities were quantitated by ImageJ. (d) The levels of TNFα in the supernatant of BMDM stimulated with fixed *C. albicans* yeast form by a MOI of 1:3. Cells were cultured in SC medium with glucose with or without lactate for overnight. *p* Values were calculated using ANOVA with Tukey post hoc analysis (\*\*\*\**p*<0.0001; *** *p*<0.001, * *p*<0.05).

Lactate is reported to reduce β-glucan exposure in yeast via stimulation of the Ace2 transcription factor (9). Since Ace2 regulates *ENG1* transcription, we examined if lactate-induced β-glucan masking is mediated by Eng1. We cultured *eng1* mutants and WT cells in the presence of lactate, fixed them, and then incubated them with BMDM. As expected, WT cells grown in lactate-containing medium induced much less TNFα secretion relative to cells grown in glucose (Fig 4d). However, *eng1* mutant cells grown in the presence of lactate stimulated significantly higher TNFα production. Therefore, lactate-induced β-glucan masking is largely dependent on Eng1. This suggests that signaling pathways that regulate β-glucan exposure may act through altering *ENG1* expression.

### The *eng1* mutant shows attenuated virulence in male mice but is hypervirulent in female mice in comparison to the wild-type strain

To evaluate the role of Eng1 during infection, we used the well-established model of hematogenously disseminated candidiasis. Male and female C57BL/6J mice were infected with 1 × 10^5^ cells of the WT, *eng1* deletion, or *ENG1* complemented strain by tail vein injection and monitored for survival. In male mice, the *eng1* mutant showed attenuated virulence. The median survival of mice infected with the WT or the *ENG1* complemented strain was 10.5-12 days, whereas the median survival of mice infected with the *eng1* cells was 19.5 days (Fig 5a, left panel). Our observation of attenuated virulence of the *eng1* mutant in male mice is consistent with the findings from two previous studies of mutants with increased β-glucan exposure in the systemic candidiasis model (12,32). The attenuated virulence is Dectin-1 dependent ((12), data not shown). Female mice were less susceptible than male mice to infection with the WT strain, as reported (33). But the *eng1* mutant was hyper-virulent in female mice. The median survival of female mice infected with the WT or *ENG1* complemented strain was more than 28 days whereas the median survival of mice infected with the *eng1* mutant yeast was 15 days (Fig 5a, right panel). The difference in survival is Dectin-1 dependent as the Dectin-1 deficient female mice showed similar susceptibility to the *eng1* deletion and *ENG1* complemented strain (Supplemental Fig. 1).

**Figure 5.**
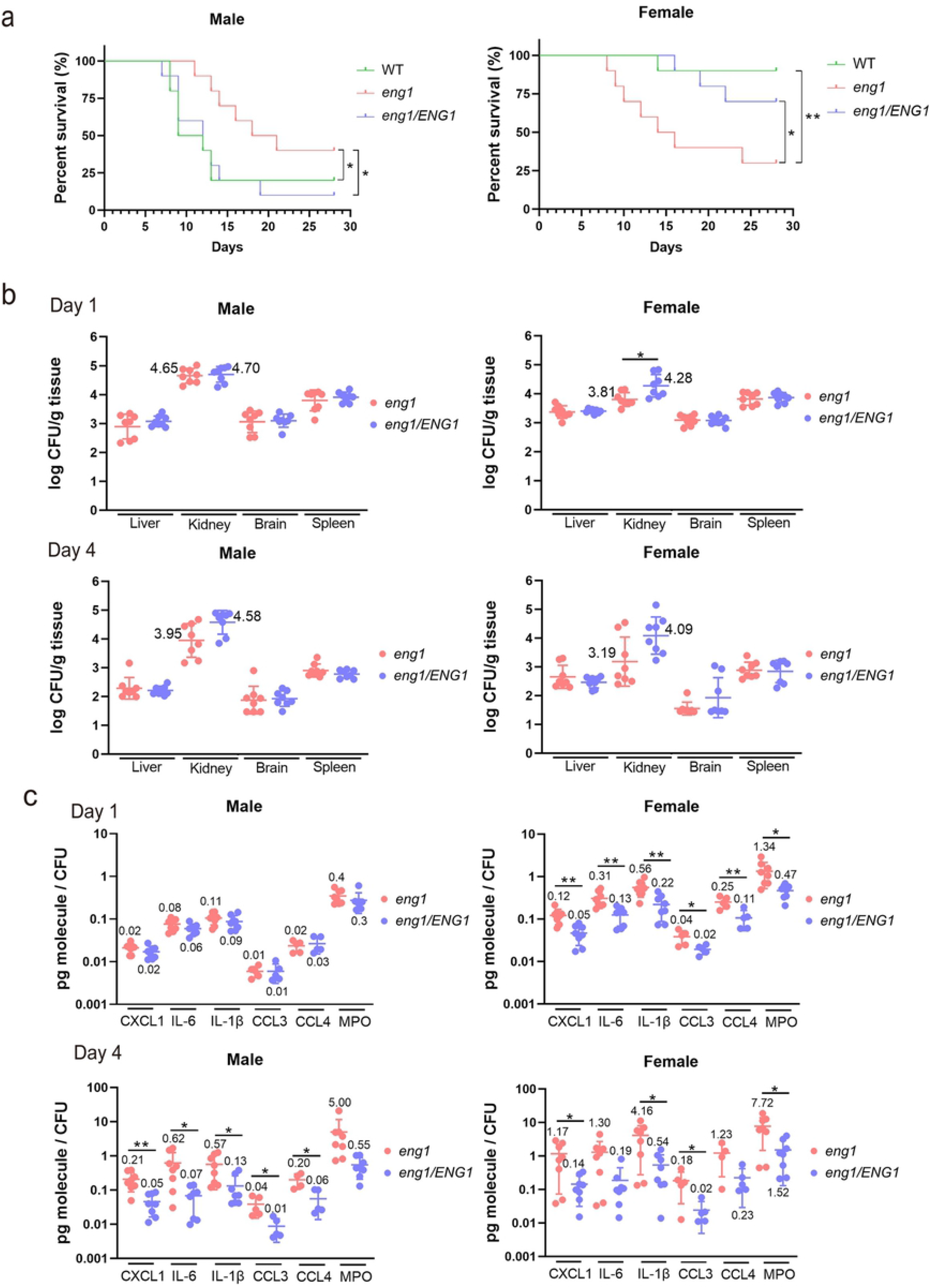
Higher immune response induced by the *eng1* mutant leads to hypo-virulence in male and hyper-virulence in female. (a) Survival of C57BL/6 male and female mice after intravenous inoculation with 1 × 10^5^ yeast phase cells of the indicated strains of *Candida albicans* (n = 10). *p* Values were calculated with Gehan-Breslow-Wilcoxon (\*\**p*<0.01; * *p*<0.05). (b) Fungal burden of the kidney, brain, spleen, and liver of mice at 1-day and 4-day postinfection after inoculation with 1 × 10^5^ *eng1* deletion mutant or *eng1/ENG1* complemented *C. albicans* yeast. Results are median ± interquartile range with 8 mice per strain. Experiments were done twice; n=3 for the first time and n=5 for the second time. *p* Values were calculated with Mann-Whitney test (\**p*<0.05). (c) Relative cytokine/chemokine or MPO levels at 1-day and 4-day post-infection. IL-1α levels were similar to IL-1β in all groups, so they were not shown in the figure. *p* Values were calculated with Unpaired t-test (\*\**p*<0.01; * *p*<0.05).

### The *eng1* mutant activates a greater renal immune response

To determine why the *eng1* mutant had different virulence in male vs. female mice, we analyzed the organ fungal burden and inflammatory response of mice after 1 and 4 days of infection. We found that the fungal burden in the liver, kidney, brain, and spleen of male mice infected with the *eng1* mutant was similar that of mice infected with the *ENG1* complemented strain at both time points (Fig. 5b). The fungal burden of the kidneys of the female mice infected with the *eng1* mutant was lower at day 1 relative to mice infected w the *ENG1* complemented strain. There was a similar trend at day 4, but the difference was not statistically significant (Fig. 5b, lower right panel).

To determine if the *eng1* mutant elicited a different inflammatory response than the *ENG1* complemented strain, we analyzed the levels of cytokines and chemokines that are up-regulated in the kidney during systemic candidiasis (4). We also measured the levels of myeloperoxidase (MPO) to assess the relative amounts of phagocytes (neutrophils, monocytes, and macrophages) (34–36) in the kidneys. We found the levels of CXCL1, IL-6, IL-1β, CCL3 and CCL4 were higher in the kidney homogenates infected with the *eng1* strain at days 1 and 4 post-infection in comparison to that infected with the *ENG1* strain (Fig 5c). Because Pearson correlation analysis indicates that cytokine levels in each kidney are highly correlated with the corresponding fungal burden (Supplemental Fig 2), the data were normalized to the kidney fungal burden of the individual mice (37). After 1 day of infection in the male mice, the chemokines, cytokines, and MPO levels were similar in animals infected with the *eng1* mutant and the *ENG1* complemented strain (Fig 5c upper panel). After 4 days infection, the levels of these inflammatory mediators in mice infected with the *eng1* mutant were significantly higher than those in mice infected with the *ENG1* complemented strain (Fig. 5c lower panel). In the female mice, the levels of chemokines, cytokines, and MPO in the kidneys were consistently higher than in the male mice (Fig 5c), indicating that female mice mounted a stronger inflammatory response to *C. albicans*. After both 1 and 4 days of infection, the levels of these inflammatory mediators were significantly higher in the mice infected with the *eng1* mutant as compared to animals infected with the *ENG1* complemented strain. Collectively, these results indicate that the *eng1* mutant induces a greater inflammatory response in the kidneys.

## Discussion

### β-glucan exposure is protected from Dectin-1 binding in yeast by Eng1 and Ywp1

β-glucan is exposed in hyphae and masked in yeast. Here, we demonstrate that in yeast, β-glucan masking is mediated by two highly expressed yeast proteins, Eng1 and Ywp1. The *C. albicans* endo-1,3-glucanase Eng1 trims excess β-glucan and reduces β-glucan exposure, similarly to HcEng1 in *H. capsulatum* (17). In *H. capsulatum*, β-glucan is also masked by an outer layer of α-glucan (38). By contrast, in *C. albicans*, β-glucan is believed to be masked by an outer mannan layer (8). Different from the current model, we find that the highly expressed yeast cell wall mannoprotein Ywp1 masks β-glucans, consistent with the finding by Granger that ectopic expression of *YWP1* in hyphae reduces β-glucan exposure (22), We revealed the Ywp1’s function in β-glucan masking by comparing Dectin-1 binding between the *eng1* and *eng1 ywp1* yeast cells. In the absence of Ywp1, Eng1 trims β-glucan not only at the septa but also over the entire cell wall. This also revealed that Eng1 not only acts at the site of septa but also over the entire cell surface when β-glucan is not masked by Ywp1. Thus, we propose that Eng1-mediated β-glucan trimming and Ywp1-mediated β-glucan masking are two parallel mechanisms utilized by *C. albicans* yeast cells to reduce β-glucan exposure (Fig 6). Ywp1 is not required for β-glucan masking at the site of septa as the *eng1* single mutant showed strong Dectin-1 binding at septa. The finding that Ywp1 reduces β-glucan exposure challenges the current model that β-glucan is protected by the outer mannan layer on the cell surface of *C. albicans* (12,39). How Ywp1 protects β-glucan exposure needs further investigation.

**Figure 6.**
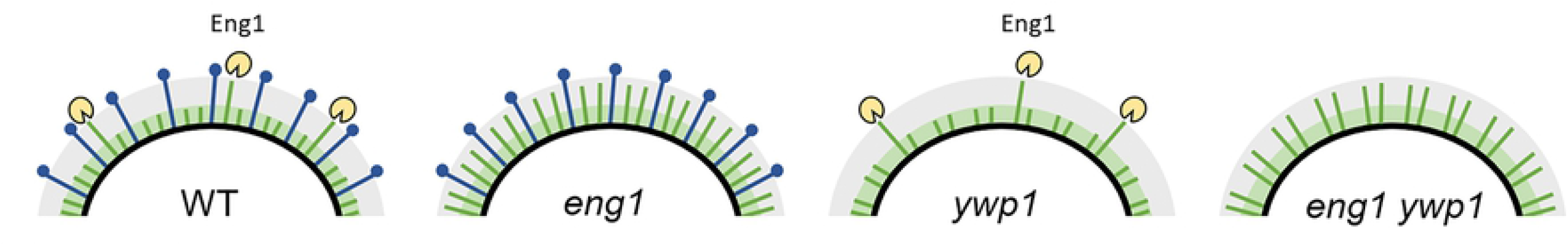
Models of β-glucan protection by Eng1-mediated trimming and Ywp1-mediated masking. Green sticks and shade represent β-glucan. Blue sticks represent Ywp1. Grey shade represents mannan layer.

Both Ywp1 and Eng1 are expressed predominately by yeast. *YWP1* is specifically expressed in yeast cells, not in hyphae. Thus, our findings directly link the dimorphic regulation of β-glucan exposure to the hyphal transcriptional program. In hyphae, phosphorylation of the Efg1 transcription factor by hypha-specific Cdk1-Hgc1 represses Ace2-regulated genes (including *ENG1*), leading to the formation of cell chains (27). In addition to being regulated by the hyphal transcriptional program, conditions that regulate cell-separation including incubation with lactate (9,16) and the anti-fungal drugs fluconazole and caspofungin also regulate *ENG1* transcription and therefore β-glucan exposure. It has been reported that Cek1 hyper-activation increases β-glucan exposure (40). This is probably caused by reduced levels of Eng1 because RNA-seq data indicate that *ENG1* transcription is repressed by the Cek1 pathway (32). The endo-1,3-β-glucanae Eng1 is likely the major glucanase for controlling β-glucan exposure based on the strong increase in the levels of cytokine production in the *eng1* mutants in comparison to the exoglucanase mutant *xog1* (16).

### β-glucan exposure in the *eng1* mutant leads to a greater immune response and modulates virulence

Pirofski and Casadevall (41) postulate that tissue damage can be caused by both the microbial pathogen and the host inflammatory response. In the current study, the kidney fungal burden of mice infected with the *eng1* mutant was generally similar to that of mice infected with the *ENG1* complemented strain, even though the survival was different. This result suggests that the altered mortality of mice infected with the *eng1* mutant was due to an alteration in the host inflammatory response elicited by greater β-glucan exposure in the *eng1* mutant. When the male mice were infected with the *eng1* mutant, the increased β-glucan exposure induced a stronger host inflammatory response, and resulted in prolonged survival of the mice. When the female mice were infected with the *eng1* mutant, the increased β-glucan exposure enhanced the already strong host inflammatory response, which caused tissue damage and resulted in reduced survival of the mice. Dectin-1 plays a protective role in defending against systemic *C. albicans* infection (6,42), as seen in the cases of female mice infected with the wild-type *C. albicans*. However, an over-exuberant immune response can also be detrimental, as seen in female mice infected with *eng1* mutant. Some of the immunopathology in the *eng1* infected mice may have been driven by high levels of CCL3 and CCL4, which can drive neutrophil-dependent immunopathogenic responses in the kidney mediated through CCR1(4,43). This study demonstrates that the level of β-glucan exposure is important for modulating the balance between immune protection and immunopathogenesis.

In summary, this study demonstrates that β-glucan exposure in yeast is protected by two highly expressed yeast proteins, the endo-1,3-β-glucanase Eng1 and the Yeast Wall Protein Ywp1. Eng1-mediated β-glucan trimming and Ywp1-mediated β-glucan masking are two parallel mechanisms utilized by *C. albicans* yeast to minimize recognition by Dectin-1. Regulating the level of β-glucan exposure in yeast is important for modulating the balance between immune protection and immunopathogenesis.

## Material and Methods

### Source of mice

C57BL/6 mice (6–10 weeks old) were from The Jackson Laboratory (Bar Harbor, ME). Dectin-1^-/-^ mice were a kind gift from Yoichiro Iwakura (University of Tokyo, Japan) bred on a BL/6 background as described (44). Animal studies were compliant with all applicable provisions established by the Animal Welfare Act and the Public Health Services Policy on the Humane Care and Use of Laboratory Animals. All animal studies were approved by the Institutional Animal Care and Use Committee (IACUC) of the Los Angeles Biomedical Research Institute and UC Irvine.

### Media and growth conditions

Strains were grown in YEP (1% yeast extract, 2% peptone, 0.015% L-tryptophan) plus 2% dextrose for yeast growth, or synthetic complete medium (0.17% Difco yeast nitrogen base w/o ammonium sulfate, 0.5% ammonium sulfate, complete supplement amino acid mixture) plus 2% N-acetylglucosamine (GlcNAc) for hyphae induction unless otherwise described. Yeast cells were cultured at 30°C. Culture for hyphae was pre-warmed and maintained at 37°C.

### Plasmid and *C. albicans* strain construction

Strains used in this study were listed in Table S1 and primers in Table S2. Gene deletion and complementation were constructed by CRISPR/Cas9 as described (28).

#### BES119-*ENG1* plasmid

The *ENG1* complementation repair template was comprised of 2 pieces of PCR products generated using primer 10,11 and primer 12,13. The repair template plasmid was constructed by integrating the PCR products into the *BES119* plasmid (45) by Gibson assembly (46) for amplification in *E. coli*. On the day of transformation, repair template plasmids were isolated with the GeneJET Plasmid Miniprep Kit (Thermo Scientific) from the overnight *E. coli* culture and digested with SacI and EcoRV

#### *eng1* deletion strain

*ENG1* was deleted in the SC5314 *LEU2* heterozygous knockout strain which was kindly provided by the Hernday lab (28). Primer 1 was used to generate the sgRNA near the 5’ end of the *ENG1* open reading frame, and primer 2 was used to generate the sgRNA near the 3’ end. Primer 3 and 4 were used for the repair template, which contained the complemented sequence of the sgRNA for *ENG1* complementation. Primer 5-9 were used to confirm the deletion of the *ENG1* DNA sequence by colony PCR.

#### *ENG1* complemented strain

*ENG1* was complemented in the *eng1* strain. Primer 9 was used to generate the sgRNA. Repair template was isolated from the BES119-*ENG1* plasmid. Primer 14 and 15 were used in couple with primer 7 and 8 to confirm the correct insertion of the *ENG1* DNA sequence by colony PCR.

#### *ywp1* and *eng1 ywp1* deletion strain

*YWP1* was deleted in the SC5314 *LEU2* heterozygous knockout strain to construct *ywp1* single deletion strain, and in *eng1* to generate *eng1 ywp1* double deletion strain. Primer 18 was used to generate the sgRNA. Primer 19 and 20 were used for the repair template. Primer 21-24 were used to confirm the deletion of the *YWP1* DNA sequence by colony PCR.

### Dectin-1-Fc staining

Staining method was derived from an established protocol (47). Soluble Fc-mDectin-1a containing the C-terminal extracellular domain of mouse Dectin-1a fused with the human IgG1 Fc domain was purchased from Invivogen. Overnight *C. albicans* yeast culture were diluted 1:20 in fresh YPD medium and grew for 6 hours for yeast form, or diluted 1:100 in fresh SC + GlcNAc and grew for 5 hours for hyphae form. 5 × 10^6^ cells were harvested, centrifuged and washed twice with PBS. After fixed with 4% formaldehyde for 15 minutes, yeast cells were washed twice with PBS and twice with binding buffer (0.5% BSA, 5mM EDTA, 2mM Sodium Azide) and incubated with 0.5 μg Dectin-1-Fc protein for 1 hour at 4 degrees. Cells were then washed twice with binding buffer and incubated with 0.5 μg secondary FITC-conjugated Rabbit anti-Human IgG for 1 hour at 4 degrees in dark. Cells were then washed three times with wash buffer and processed for microscopy. Images were obtained on a Zeiss Axioplan 2 or an inverted Zeiss Axio Observer.Z1 microscope (Carl Zeiss MicroImaging, Inc., Thornwood, NY) fluorescent system equipped with the AttoArc HBO 100 and the X-Cite series 120 mercury lamps, respectively. Both fluorescence microscopes were equipped with GFP and DAPI (4′,6′-diamidino-2-phenylindole) filter sets. Images were taken using a ×100 NA 1.4 objective lens. Processing was done using the software ImageJ (National Institutes of Health, USA), as well as Photoshop and Illustrator (Adobe Systems, Inc., Mountain View, CA). Mean fluorescence intensity was calculated based on at least three randomly selected images by ImageJ. MFI values were formulated as the average integrated intensity of the fluorescence subtracted by background signal.

### Macrophage infection and cytokine measurement

BMDM derivation was carried out as described (48). Bone marrow cells were harvested from the femurs and tibia of C57BL/6 WT or Dectin-1^-/-^ mice. The bones were trimmed at each end and centrifuged at 800 × g for 10 s, and bone marrow cells were suspended in growth medium (Dulbecco’s modified Eagle’s medium plus10% FBS and 1% penicillin/streptomycin). Macrophages were derived by culturing bone marrow cells in growth medium plus 20ng/mL mCSF for 7 days. Medium with fresh mCSF was added every 2-3 days. Then macrophages were harvested by treating with Cell Stripper solution (Corning) for 15 minutes and scraping. Cells were counted with a hemocytometer and 10^5^ cells were plated to each well in a 96 well plate for overnight. Overnight *C. albicans* yeast culture were diluted 1:20 in fresh YPD medium and grew for 6 hours. Cells were then fixed with 4% formaldehyde and washed 4 times with PBS. Bone marrow-derived macrophages were stimulated with fixed yeast (MOI=3) or live overnight yeast (MOI=1) for 6 hours at 37 degrees with 5% CO_2_. Co-culture supernatant was collected and cytokines were measured by commercially available Ready-Set-Go cytokine kits (eBioscience).

### Quantitative PCR

Methods for RNA isolation were carried out as previously described (49). cDNA was synthesized from 1μg total RNA using the BioRad iScript Reverse Transcription Kit. Quantitative PCR using the BioRad SYBR Green mix and primer 16 and 17 was performed on the BioRad iCycler. Cycle conditions were 95°C for 1min, then 39 cycles of 95°C for 10s, 56°C for 45s, and 68°C for 20s.

### *In vivo* assays

*In vivo* assays were carried out as described (50). Male and female C57BL/6 and Dectin-1^-/-^ mice (6-8 weeks) were inoculated via the lateral tail vein with 1×10^5^ *C. albicans* cells per mouse. They were monitored 3 times daily for survival. For fungal burden and cytokine measurements, the mice were sacrificed after 1 and 4 days of infection. One kidney from each mouse was harvested, weighed, and homogenized. An aliquot of the homogenate was quantitatively cultured and the remaining sample was clarified by centrifugation. The supernatant was collected and stored at −80°C. On a later date, the cytokine content of the homogenates was determined by Luminex cytometric bead array.

### Statistical Analysis

At least three biological replicates were performed for all experiments, and the results are expressed as mean values ± standard deviation. Data were analyzed using student t-test, ANOVA with Tukey post hoc or Mann-Whitney test by GraphPad Prism (ver. 8.0) as indicated in the figure legends. A probability level of 5% (p < 0.05) was considered significant. Pearson correlation analysis was done by R Studio.

## ACKNOWLEDGEMENTS

The authors would like to thank Dr. Hernday for *C. albicans* strains and CRISPR plasmids. This work was supported in part by the National Institute of General Medical Sciences R01GM117111 (HL), the National Institute of Allergy and Infectious Diseases R01AI124566 (SGF), the National Institute of Dental and Craniofacial Research R01DE026600 (SGF), as well as by the National Eye Institute R01EY18612 (EP).

---

Supplemental Figure 1. Survival of C57BL/6 control and Dectin-1^-/-^ female mice after intravenous inoculation with 1 × 10^5^ yeast phase cells of the indicated strains of *Candida albicans* (n = 5 for B6 and n=3 for Dectin-1^-/-^ mice).

Supplemental Figure 2. Pearson correlation analysis of cytokine/chemokine levels and fungal burdens.

